# Mitochondrial and Cardiolipin Adaptations to Ventricular Assist Device Support in Pediatric Versus Adult Failing Myocardium

**DOI:** 10.64898/2026.04.01.715996

**Authors:** Caitlyn S. Conard, Mariana Casa de Vito, Obed O. Nyarko, Raleigh L. Jonscher, Elisabeth K. Phillips, Kathryn C. Chatfield, Amrut V. Ambardekar, Jordan Hoffman, Scott R. Auerbach, Matthew L. Stone, Carmen C. Sucharov, Brian L. Stauffer, Genevieve C. Sparagna, Shelley D. Miyamoto

**Affiliations:** Department of Medicine, Division of Cardiology, University of Colorado Anschutz Medical Campus, Aurora, Colorado USA; Department of Pediatrics, Division of Cardiology, University of Colorado Anschutz Medical Campus and Children’s Hospital Colorado, Aurora, Colorado USA; Department of Surgery, Division of Cardiothoracic Surgery, University of Colorado Anschutz Medical Campus and Children’s Hospital Colorado, Aurora, Colorado USA; Division of Cardiology, Denver Health Medical Center, Denver, Colorado USA

## Abstract

**Background:** Ventricular assist devices (VADs) are used as treatment for end-stage heart failure in children and adults. We previously demonstrated decreased mitochondrial function and changes in cardiolipin, a mitochondrial phospholipid, in explanted pediatric and adult failing hearts. In this study, we tested the hypothesis that VAD unloading of failing hearts leads to positive changes in myocardial cardiolipin in both pediatric and adult hearts.

**Methods:** Ventricular tissue was collected from the same patient at time of VAD implantation and at transplant. Ejection fraction (EF), left ventricular internal diameter at end-diastole (LVIDd) and brain natriuretic peptide (BNP) were assessed pre- and post-VAD. Cardiolipin species from paired VAD core and explants were quantified using liquid chromatography mass spectrometry. Mitochondrial respiration was measured in ventricular tissue pre- and post-VAD in paired pediatric samples using the Oroboros Oxygraph-2k.

**Results:** VAD support led to increased EF and decreased LVIDd and BNP. The predominant cardiolipin species in cardiac mitochondria, tetralinoleoylcardiolipin, was positively remodeled in pediatric post-VAD myocardium, while adult post-VAD myocardium demonstrated significantly increased total cardiolipin and decreased oxidized cardiolipin but did not demonstrate the tetralinoleoylcardiolipin remodeling seen in pediatric hearts. In pediatric patients, VAD support resulted in significant increases in Complex I+II activity, and a trend toward increases in Complex I activity.

**Conclusion:** Our data demonstrate age-related differences in VAD-associated cardiolipin remodeling and suggest that improved mitochondrial function in pediatric VAD-supported hearts could be related to increased tetralinoleoylcardiolipin.

## Introduction

Despite advancements in medical therapy for the treatment of heart failure (HF), some patients require ventricular assist device (VAD) or heart transplantation (1). In both children and adults, mechanical unloading with a VAD is used as a bridge to heart transplant or as a bridge to recovery (2). Among other adaptations, VAD support results in reverse remodeling of ventricular size and function (3), while at the cellular level, VAD support has been shown to result in gene expression changes (4), recovery of dystrophin protein (5), resolution of cardiomyocyte hypertrophy (6) and improvement in calcium handling (7,8). While there are mixed results regarding the impact of oxidative function of cardiac mitochondria in adults following mechanical unloading with a VAD (9), the impact of VAD support on cardiac mitochondrial function in children is unknown.

Altered mitochondrial bioenergetics and ATP supply have long been recognized as a central contributor to HF progression(10). The failing heart has been shown to be functionally deficient in energy production with a switch away from fatty acid beta-oxidation (FAO), impaired oxidative phosphorylation, reduced ATP production, and increased reactive oxygen species generation(11). Cardiolipin (CL) is a mitochondria-specific phospholipid having four fatty acyl chains that is predominately localized to the inner mitochondrial membrane(12). CL is essential for the structural integrity and optimal function of most electron transport chain (ETC) proteins and supports the activity of numerous other mitochondrial proteins that depend on it to function(13). In the healthy heart, CL exists predominantly as tetralinoleoylcardiolipin (L_4_CL), having four linoleate (LA, 18:2n6) side chains. We have shown that LA side chains on CL promote FAO and that the substitution of oleate (OA, 18:1n9) for LA results in substrate switching away from FAO(14). In healthy heart mitochondria, CL is remodeled from its nascent form to its predominant L_4_CL form. This remodeling in cardiac mitochondria is primarily through two alternative enzymatic mechanisms. In the first, a transacylase, tafazzin, exchanges the nascent fatty acyl chain for LA. The second mechanism involves a two-step process where phospholipase A_2_ (PLA_2_) cleaves an acyl group forming monolysocardiolipin (MLCL; CL with 3 acyl chains). MLCL can then be remodeled to L_4_CL when the enzyme MLCL acyl transferase (MLCLAT) adds an LA back onto MLCL and repeats this process until all the side chains are LA (14,15). In HF, CL composition becomes pathologically remodeled, with depletion of L_4_CL and accumulation of more saturated species, such as CL with side chains consisting of OA, impairing mitochondrial respiration and FAO(17,18).

We previously demonstrated diminished mitochondrial respiration and abnormal CL remodeling in pediatric and adult failing myocardium (17,18). However, whether mechanical unloading due to VAD support in pediatric patients reverses these mitochondrial abnormalities in CL and improves mitochondrial function, and whether CL changes differ between pediatric and adult patients remain unknown. The objective of this study was to use paired myocardial samples pre- and post-VAD in children and adults to define CL remodeling in unloaded pediatric and adult hearts, investigate whether VAD associated CL changes differ between these two populations, and determine if mechanical unloading is associated with improvements in mitochondrial function in children.

## Methods

### Study Population

Pediatric and adult patients with advanced HF undergoing VAD implantation as a bridge to heart transplantation were included. Systemic ventricular tissue was collected at VAD implantation (VAD core) and again at the time of transplantation (explant). Pediatric and adult cohorts were analyzed separately. The pediatric participants were females and males younger than 18 years of age with HF secondary to various forms of cardiomyopathy or congenital heart disease. The adult participants were females and males ≥18 years of age with HF secondary to dilated cardiomyopathy (idiopathic, ischemic, or familial). All races and ethnic backgrounds were included in the study. More details are available in supplemental table S-1. Hearts were donated by patients or their families to the University of Colorado Institutional Review Board (IRB)-approved pediatric and adult cardiac biobank.

### Tissue Collection

Left ventricular (or systemic ventricle in the case of children with single ventricle congenital heart disease) tissue was obtained during VAD implantation (apical core) and at the time of heart transplantation. Fresh ventricular tissue was quickly dissected in cold Tyrode’s solution in the operating room at time of collection. Pediatric samples designated for mitochondrial respiration were freshly stored in BIOPS solution (19) and mitochondrial respiration studies performed within 24 hours. Parallel samples were flash frozen at time of collection and stored at −80°C for CL analysis.

### Clinical and Echocardiographic Data

Echocardiographic data including, ejection fraction (EF) and left ventricular internal diameter in cm at end-diastole (LVIDd) were determined based on review of clinical echocardiogram reports within three weeks prior to VAD implantation and within three weeks prior to transplantation. Pediatric patients excluded from the Echo portion of the study included patients with a single ventricle (n=2), 1 patient that was on a VAD for only 2 days, and 1 patient that had cardiac surgery and was on extracorporeal membrane oxygenation (ECMO) prior to VAD placement. BNP levels were obtained from the medical record for the same time points. BNP levels exceeding laboratory detection limits (5000 ng/L or 2500 ng/L) were reported at those limits for recording purposes. More details are available in supplemental table S-2.

### Cardiolipin Quantification

Frozen tissue was homogenized in PBS. Lipids were extracted as previously described using a modified Bligh and Dyer protocol with 100 nmol 1,1′,2,2′-tetramyristoyl CL used as an internal standard (Avanti Polar Lipids, Alabaster, AL US). Normal phase high pressure liquid chromatography coupled to electrospray ionization mass spectrometry (LC-ESI-MS) was performed in the Sciex API4000 as previously described(14). Standard curves were made using tetraoleoylcardiolipin (Avanti Polar Lipids, Alabaster, AL US) as a reference standard.

### Cardiolipin Data Presentation

Fifteen separate CL molecular species were measured both in absolute amounts of nmol/mg protein and in the percentage of each species comprising the CL profile. This profile was mathematically determined by dividing each molecular species by the sum of all 15 species. More CL species details can be found in supplemental table S-3 and supplemental table S-4. The percentage allows insight into molecular species remodeling. CL has 4 fatty acyl chains which in the healthy heart are predominately LA, but are often substituted with OA and other fatty acyl chains in HF in adult and pediatric patients(17,18). Four mathematical constructs were calculated to give a snapshot of CL alterations: (1)**Total CL** is the sum the 15 detected CL species; (2)**L**_**4**_**CL/L**_**3**_**OCL**: L_4_CL has 4 LA side chains having a mass to charge ratio (m/z) of 1448 and a carbon to double bond ratio (C:DB) of 72:8. L_3_OCL is CL which has 3 LAs and one OA (m/z 1450; C:DB 72:7). This ratio indicates the degree of CL remodeling; (3) **L**_**3**_**MLCL/Total CL**: MLCL with three LA side chains (L_3_MLCL; m/z 1186; C:DB 54:6) was divided by total CL to show the degree of turnover of CL; (4) **L**_**4**_**CLox/L**_**4**_**CL**: Oxidized L_4_CL (L_4_CLox) was detected as m/z 1464 (1448+O16). It is expressed as a fraction of its non-oxidized parent L_4_CL.

### Mitochondrial Respiration

Respiration of permeabilized cardiac fibers, stored in BIOPS (20) as described above, was measured by high-resolution respirometry (Oxygraph-2k, Oroboros Instruments, Innsbruck, Austria) using a series of substrates in a stepwise protocol to evaluate various components of the electron transport chain. Specifically, tissue was cut into approximately 2-mg pieces and teased using forceps to separate fibers. The tissue was then placed in a solution of BIOPS containing 30 μg/ml saponin for 30 min to permeabilize the plasma membrane and to allow substrate delivery to the mitochondria. Fibers were washed for 10 min at 4°C in ice-cold mitochondrial respiration medium (Mir05) containing 25 μM blebbistatin(19). Samples were blotted on filter paper and weighed. Mitochondrial function was measured in the Oroboros O2K at 37°C containing MirO5 plus 25 μM blebbistatin. Complex function was measured by stepwise addition of 5 mM pyruvate, 1 mM malate, 4 mM adenosine diphosphate (ADP), 10 mM glutamate, and 10 mM succinate. Oxygen flux rates were normalized per milligram of wet tissue weight as described in our previous publication(21). These experiments were performed only in pediatric hearts.

### Statistical Analysis

Paired analyses were performed comparing pre- and post-VAD samples within each age group using Prism v. 11(22). Statistical significance was defined as p<0.05 and trends expressed for p<0.1. Statistical tests are listed in the figure legend. Outliers were calculated using two standard deviations above or below the mean. Unless otherwise stated, data are expressed as mean ± standard deviation.

## Results

### Patient Characteristics

The summary of the subjects’ clinical characteristics and demographics are found in Table 1. There were 31 adult HF patients (median age of 55.8 years and IQR 13.8 years, mean days on VAD 266 ± 272.7 and IQR of 223, 90.3% males, 23 on a HeartMate II, 8 on a HeartWare HVAD).

**Table 1.** The summary of the subjects’ clinical characteristics and demographics.

|  | Number of Patients (N), % males | Median Age, (IQR*) | Average Days on VAD*, SD*, (IQR) | HeartMate III VAD | HeartMate II VAD | HeartWare Ventricular Assist System (HVAD) | Berlin Heart EXCOR |
| --- | --- | --- | --- | --- | --- | --- | --- |
| Adult | 31, 90.3 | 55.8, (13.8) | 265.8, 272.7, (223) | 0 | 23 | 8 | 0 |
| Pediatric | 22, 63.6 | 10.16, (11.4) | 76.1, 99.8, (75.25) | 5 | 0 | 7 | 10 (2 with BiVAD* support) |

\*Abbreviations: Interquartile Range (IQR); Ventricular Assist Device (VAD); Standard Deviation (SD); Biventricular assist device (BiVAD)

There were 22 pediatric patients included in this study (median age of 10.16 years and IQR 11.4 years, mean days on VAD 76 ± 99.8 and IQR of 75.25, 63.6% males, 10 on a Berlin Heart EXCOR (2 with a biventricular assist device), 7 on a HeartWare (HVAD), and 5 on a HeartMate III). More details are available in supplemental table S-1.

### VAD Support Leads to Reverse Remodeling in Both Pediatric and Adult Patients

We assessed the change in echocardiographic parameters and BNP in pediatric and adult patients supported with a VAD (Fig.1). Both children and adults demonstrated improvements in systolic function and ventricular dimensions following VAD support. VAD support resulted in a significant increase in EF (n=18 pediatric patients, pre VAD EF 20.29 ± 10.8 and post VAD EF 31.28 ± 15.46, p=0.0190; n=17 adults, pre VAD EF 15.51 ± 7.82 and post VAD EF 35.75 ± 18.56, p=0.0002, Fig. 1A), significant decrease in LVIDd (n=16 pediatric patients, pre VAD LVIDd 5.82cm ± 1.62 and post VAD LVIDd 4.94cm ± 1.48, p<0.0001; n=24 adults, pre VAD LVIDd 6.91cm ± 1.29 and post VAD LVIDd 5.07cm + 1.63, p<0.0001, Fig. 1B), and significantly decreased BNP levels (n=15 pediatric patients, pre VAD BNP 2255 ± 1494 and post VAD BNP 418.1 ± 343.7, p=0.0004; n=24 adults, pre VAD BNP 1554 ± 826.3 and post VAD BNP 260.3 ± 200.4, p<0.0001, Fig. 1C). More details are available in supplemental table S-2.

**Figure 1:**
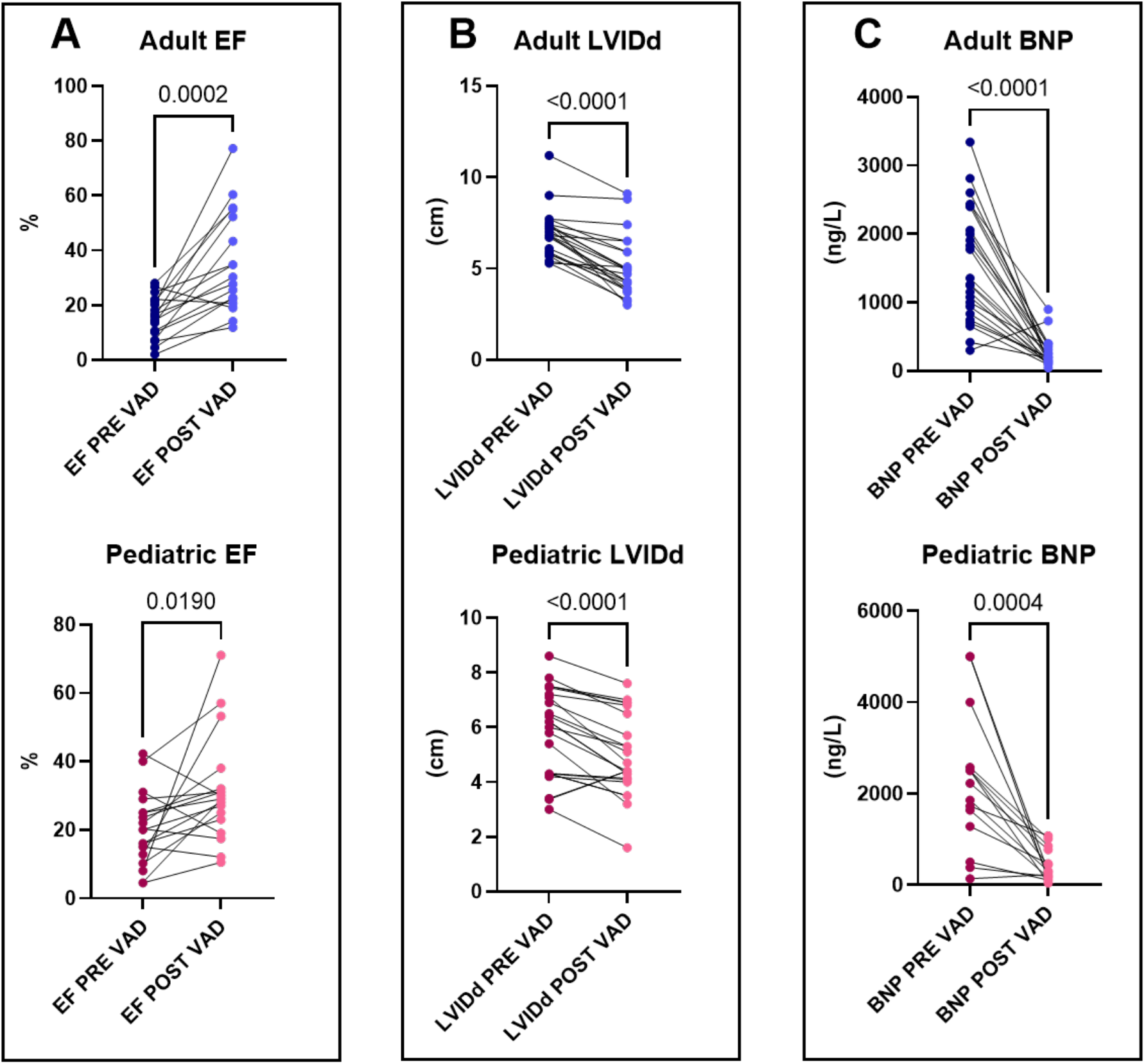
Clinical Reverse Remodeling in Adult and Pediatric Heart Failure Following VAD Support. Mechanical unloading with a ventricular assist device (VAD) induces significant reverse remodeling in both adult and pediatric cohorts. Echocardiographic and clinical parameters were measured pre- and post-VAD, demonstrating improvements in systolic function and reduced BNP. **(A)** Ejection fraction (EF) significantly improved in both adults and pediatrics n=17 adults, n=18 pediatric. **(B)** Left ventricular internal diameter at end-diastole (LVIDd) significantly decreased post-VAD n=24 adults, n=16 pediatric. **(C)** Brain natriuretic peptide (BNP) levels, a marker of cardiac strain, were significantly reduced in both populations n=24 adults, n=15 pediatric. Paired analyses were performed comparing pre- and post-VAD samples within each age group. Number above each graph shows p-value.

### Pediatric Hearts Exhibit Selective Cardiolipin Remodeling

We analyzed 15 different CL species in paired VAD core and explanted myocardium. There were no significant differences in the sum of CL species (Total CL) in the pediatric population between the VAD core and the explanted myocardium (n=19, pre-VAD 8.87 ± 3.89 nmol/mg and post VAD 10.27 ± 3.74 nmol/mg, p=0.2017, Fig.2A). Following VAD support, the pediatric explanted myocardium demonstrated a significant increase in the L_4_CL/L_3_OCL ratio (n=20, pre-VAD 3.83 ± 0.71 and post-VAD 4.35 ± 1.13, p=0.0444, Fig.2B) and an increased L_3_MLCL/Total CL ratio compared to the VAD core (n=20, pre VAD 0.022 ± 0.0056 and post VAD 0.029 ± 0.0085, p=0.0098, Fig. 2C). There was no significant change in the ratio of oxidized L_4_CLox/L_4_CL between VAD core and explanted myocardium (n=20, pre-VAD 0.004 ± 0.003 and post VAD 0.0033 ± 0.0027, p=0.2170, Fig. 2D).

**Figure 2:**
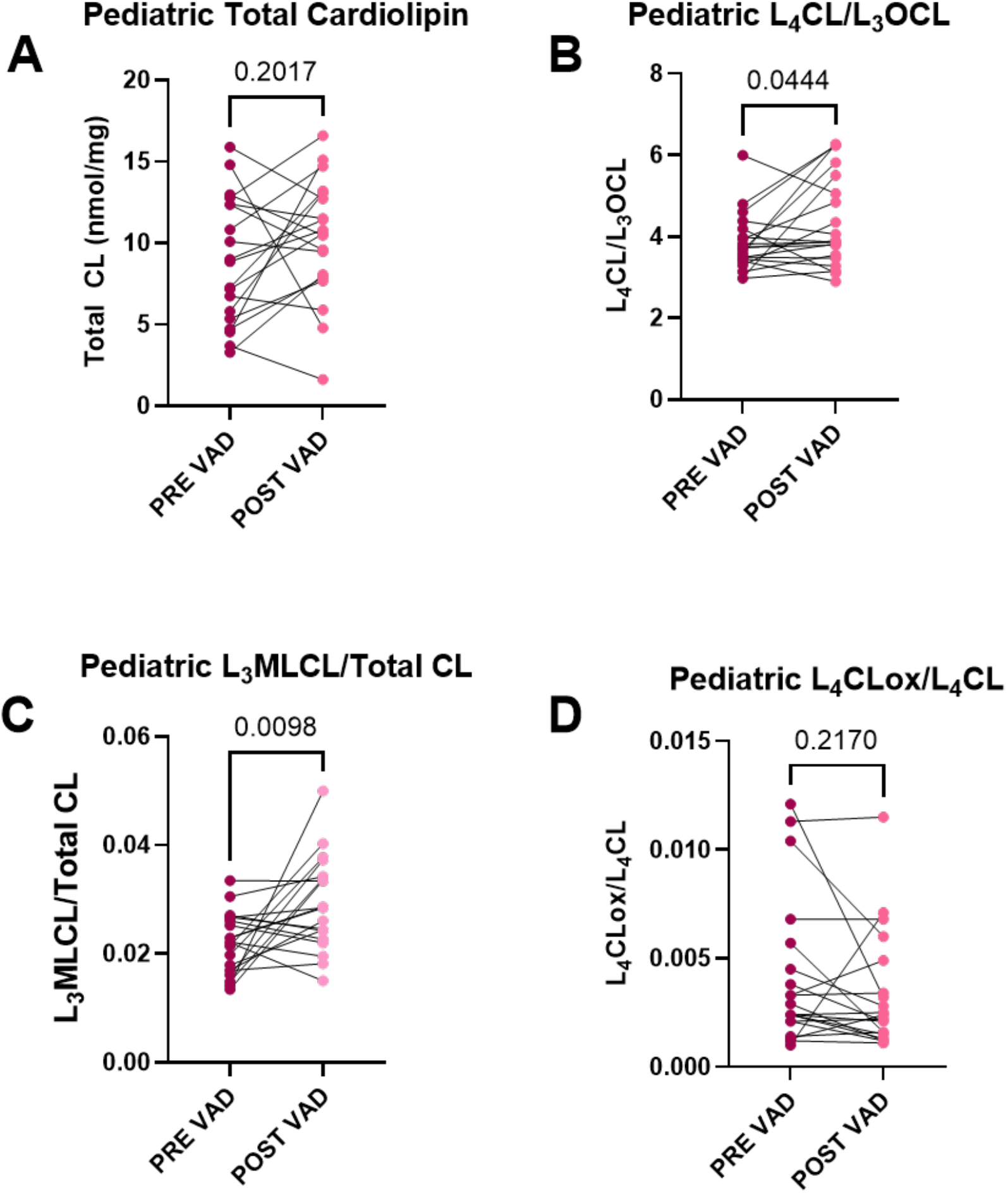
Pediatric Hearts Exhibit Beneficial Remodeling of Cardiolipin: In pediatric hearts pre-vs post-VAD **(A)** There was no significant change in total CL content n=19 pairs; **(B)** the ratio of L_4_CL to L_3_OCL, n=20 pairs, significantly increased; **(C)** the monolysocardiolipin ratio (L_3_MLCL/Total CL) significant increased, n=20 pairs; **(D)** there were no significant changes in the oxidized L_4_CLox/L_4_CL ratio, n=20 pairs. The number above each graph shows p-value.

### Adult Hearts Demonstrate Distinct Cardiolipin Adaptations

In contrast to what was seen in the pediatric heart, adult explanted myocardium following VAD support exhibited a significant increase in total CL (n=31, pre-VAD 10.82 ± 6.46 nmol/mg and post-VAD 15.08 ± 6.29 nmol/mg, p=0.0060, Fig. 3A). Unlike in pediatric myocardium, in VAD-supported adult myocardium there was no significant increase in remodeling of L_4_CL, expressed as the L_4_CL/L_3_OCL ratio (n=31, pre-VAD 3.32 ± 0.9258 and post-VAD 3.66 ± 0.8089, p=0.1473, Fig. 3B) and no change in the L_3_MLCL/Total CL ratio (n=30, pre-VAD 0.040 ± 0.0187 and post-VAD 0.0351 ± 0.017, p=0.3147, Fig. 3C). However, VAD support in adults resulted in decreased levels of the oxidized CL ratio, L_4_CLox/L_4_CL in the explanted myocardium (n=30, pre-VAD 0.0064 ± 0.0077 and post VAD 0.0028 ± 0.00189, p=0.0262, Fig. 3D).

**Figure 3:**
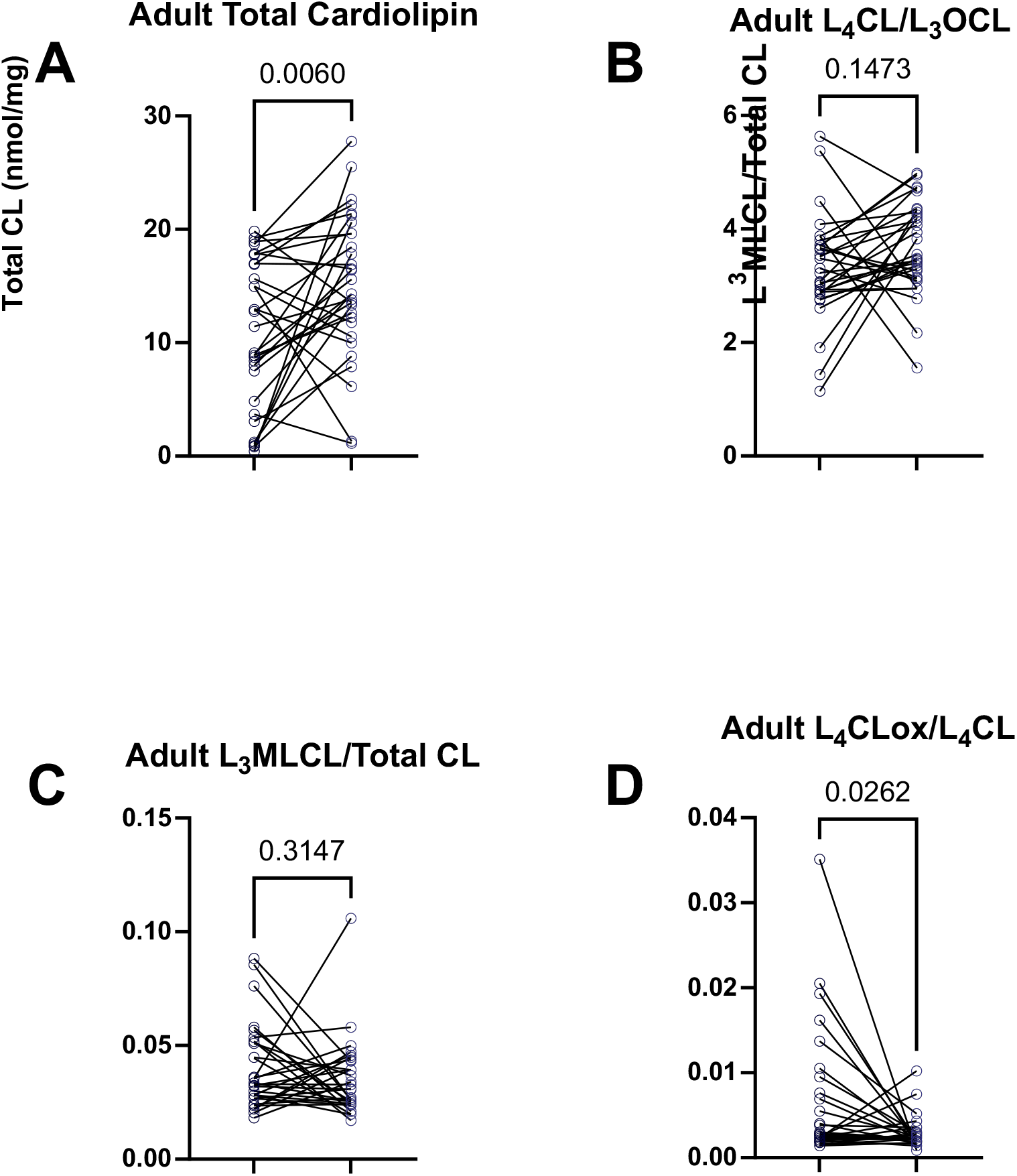
Adult Hearts Show an Increase in Total and Oxidized Cardiolipin: In adult hearts pre-vs post-VAD **(A)** Total CL content was significantly increased, n=31 pairs; **(B)** the remodeling ratio of L_4_CL to L_3_OCL was unchanged, n=31 pairs; **(C)** the monolysocardiolipin ratio (L_3_MLCL/Total CL) was unchanged, n=30 pairs; **(D)** Oxidized L_4_CLox/L_4_CL was significantly decreased, n=30 pairs. The number above each graph shows p-value.

### VAD Support Improves Mitochondrial Respiration in Pediatric Hearts

To assess mitochondrial respiration in pediatric hearts, we evaluated mitochondrial function from paired freshly procured VAD core and explanted myocardium. Mechanical unloading was associated with enhanced mitochondrial respiratory capacity in pediatric myocardium. Specifically, there was a trend toward increased Complex I activity (n=6, pre VAD 47.15 ± 12.07 pmol/(s*mg) and post VAD 62.60 ± 13.72 pmol/(s*mg), p=0.0562, Fig. 4A) and a significant increase in Complex I+II activity (n=6, pre VAD 85.88 ± 19.64 pmol/(s*mg) and post VAD 102.9 ± 14.14 pmol/(s*mg), p=0.0272, Fig.4B).

**Figure 4:**
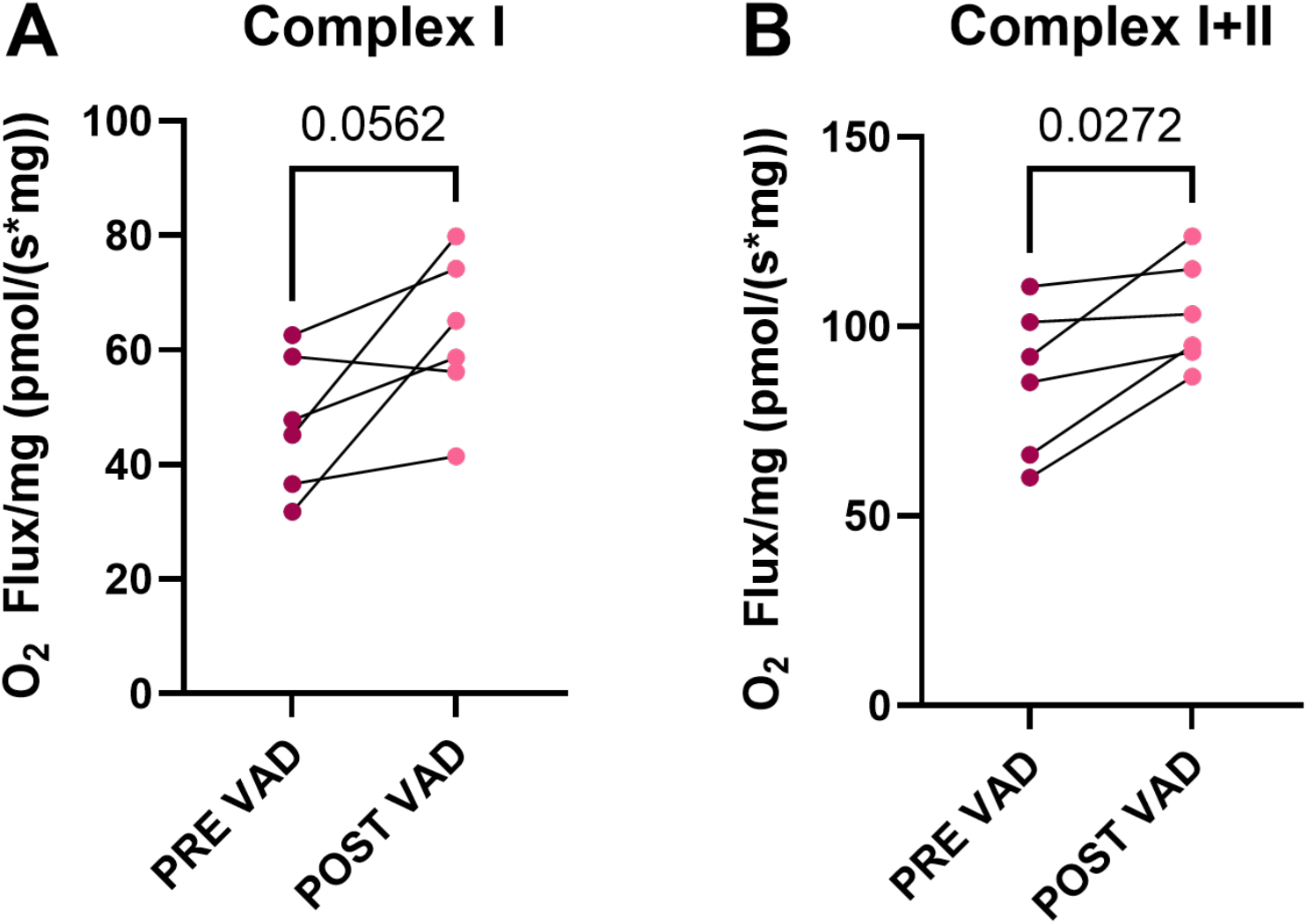
VAD Support Enhances Mitochondrial Respiration in Pediatric Myocardium. High-resolution respirometry was performed on permeabilized cardiac fibers from fresh pediatric ventricular tissue pairs (n=6) to assess mitochondrial oxidative capacity pre- and post-VAD. **(A)** Complex I–supported respiration (evaluated using malate, pyruvate and glutamate) demonstrated a trend toward an increase following mechanical unloading. **(B)** Complex I+II–supported respiration (evaluated using the further addition of succinate) increased significantly post-VAD. Oxygen flux rates are normalized per milligram of wet tissue weight. Number above each graph shows p-value.

## Discussion

This study demonstrates that while mechanical unloading with VAD support results in reverse cardiac remodeling by echocardiogram and an improvement in BNP in both pediatric and adult HF patients, the underlying mitochondrial CL adaptations differ fundamentally by age. Our findings build upon the 2002 study by the Schlame laboratory which showed that mechanical unloading by VADs promoted positive CL remodeling(23). However, this study was performed only in adults, in pre- and post-VAD hearts that were not paired, and was limited to investigation of only 3 CL species(15,23). Our study utilizes tissue from the same patient before and after VAD therapy and investigated a more comprehensive panel of 15 CL species in both pediatric and adult patients. More details on CL species can be found in supplemental table S-3 and supplemental table S-4.

VAD support in pediatric hearts led to an increase in the L_4_CL/L_3_OCL ratio without increasing total CL content. We previously showed that higher LA-containing CL species and less OA-containing species are likely associated with a shift to more FAO(24). The heart under normal conditions relies on FAO to produce most of its energy, but in HF, there is a switch away from FAO(25). Here we show that mechanical unloading of the pediatric heart resulted in a shift in the composition of CL that would favor FAO for energy generation. The increase in the L_3_MLCL/Total CL ratio seen in the pediatric hearts also suggests that VAD support of pediatric failing hearts may increase the CL turnover rate. This adaptation could indicate that volume unloading of pediatric hearts promotes positive CL remodeling via the MLCLAT pathway.

In contrast, the CL remodeling pattern in adult VAD supported hearts was distinct from what was shown to occur in the pediatric hearts. Adult hearts supported by a VAD demonstrated a significant increase in total CL, which could either represent an increase in the CL contained within each mitochondrion or perhaps an increased number of total mitochondria. Adult unloaded hearts also showed a decrease in oxidized CL whereas pediatric hearts had no change in oxidized CL. These differences in CL remodeling between pediatric and adult VAD-supported hearts may indicate that while in pediatric hearts, VADs result in a more functional remodeling pattern of CL that would be predicted to increase FAO, the adult VAD supported hearts compensate through an increase in CL or mitochondrial number without an expected impact on substrate switching. Further, because oxidized cardiolipin serves as an apoptotic signal when it translocates to the outer mitochondrial membrane(26), the increase in CL content combined with the decrease in oxidized CL in adult VAD supported hearts could reflect a decrease in the rate of apoptosis resulting in an increase in mitochondrial number.

One study in adults demonstrated that VAD support resulted in an increase in mitochondrial function in Complex I (glutamate as a substrate), but not Complex I+II(succinate as a substrate)(9). In pediatric patients using paired myocardial samples, we show that VAD support resulted in significant increases in Complex I+II activity and a trend toward improved Complex I activity. Mechanistically, the age-related variability in mitochondrial functional recovery may reflect the differences in cardiolipin remodeling between pediatric and adult VAD supported hearts, particularly given Complex II’s dependence on cardiolipin(27).

### Translational and Clinical Implications

The identification of age-specific CL plasticity is intriguing and could help inform the potential for ventricular recovery in children supported with a VAD. While VAD explant after ventricular recovery or remission is uncommon in children, there is increased interest in identifying characteristics of children that could be candidates for VAD explant without the need for a heart transplant (28). For example, if the circulating L_4_CL/L_3_OCL ratio is a surrogate for the myocardial ratio, this could be assessed in children with HF using a blood spot test or in white blood cells. This ratio could then serve as a highly specific, novel biomarker for assessing favorable myocardial metabolic adaptations in children with HF either in response to medical therapy or a VAD.

Furthermore, these distinct biochemical responses represent an opportunity for age-specific mitochondrial therapies to optimize VAD mediated recovery. For adult patients, whose recovery can be hampered by excessive mitophagy mediated by lipid oxidation signaling, adjunctive therapies utilizing targeted antioxidants could reduce CL oxidation and augment mitochondrial number during heart failure. Conversely, therapeutic strategies, such as dietary LA supplementation, designed to promote endogenous CL remodeling pathways are promising for maximizing the innate recovery mechanisms that appear to be uniquely present in pediatric hearts(14).

## Limitations and Future Directions

This study has several notable limitations. Findings are limited by the varied etiologies of HF in the included patients, the male predominance (especially in the adult cohort), and in the variation in VAD support which included both pulsatile (Berlin Heart EXCOR) and continuous flow devices (HeartMate II and III, and HeartWare). The availability of fresh tissue for high-resolution mitochondrial functional assays was limited to a modest sample size within only the pediatric cohort.

Future investigations must build upon these findings by delineating the precise enzymatic and molecular pathways responsible for the selective CL remodeling observed in unloaded pediatric myocardium by studying how FAO, mitochondrial number, and apoptosis differ between the pediatric and adult failing heart. Investigating paired circulating and myocardial L_4_CL/L_3_OCL ratios in children with HF are needed to determine if this ratio could serve as a novel biomarker of favorable lipid remodeling in children with heart failure. Importantly, determining whether dormant CL restorative pathways can be pharmacologically reactivated in the adult myocardium will be a crucial next step in developing targeted therapies to enhance myocardial recovery across all ages.

## Conclusion

Mechanical unloading with VAD support induces age-dependent mitochondrial CL remodeling. While pediatric hearts demonstrate significant CL plasticity, resulting in the selective restoration of LA-rich CL and improved fatty acid oxidation capacity, adult VAD supported hearts have higher total CL and lower oxidized CL, which could reflect an alternate mechanism for enhancing energetics. These findings position CL remodeling as a possible mechanistic target, especially for pediatric myocardial recovery.

## Acknowledgments

The University of Colorado human tissue bank is supported by NIH/NCATS Colorado CTSA Grant Number UM1 TR004300. Contents are the authors’ sole responsibility and do not necessarily represent NIH views. We would like to acknowledge the Heart Transplant Team at Children’s Hospital Colorado and UC Health University of Colorado Hospital.

